# TDP-43 regulates the expression levels of Disc-large in skeletal muscles to promote the assemble of the neuromuscular synapses in Drosophila

**DOI:** 10.1101/716423

**Authors:** Nina Strah, Giulia Romano, Clelia Introna, Raffaella Klima, Aram Megighian, Monica Nizzardo, Fabian Feiguin

**Affiliations:** International Centre for Genetic Engineering and Biotechnology, Padriciano 99, 34149 Trieste, Italy; Department of Biomedical Sciences, University of Padova, via Marzolo 3, 35131 Padova, Italy; Department of Pathophysiology and Transplantation (DePT), Dino Ferrari Centre, University of Milan, Neuroscience Section, IRCCSFoundation Ca’ Granda Ospedale Maggiore Policlinico, Via Francesco Sforza 35, 20122, Milan, Italy

**Keywords:** TDP-43, Skeletal muscles, Dlg, neuromuscular junctions, ALS

## Abstract

**Background:** Alterations in the intracellular distribution of TDP-43 were observed in the skeletal muscles of patients suffering from ALS. However, it is not clear whether these modifications play an active role in the disease or represent a physiological adaptation to muscles homeostasis.

**Result:** To answer these questions, we modulated the activity of this protein in Drosophila muscles and observed that TDP-43 was required in these tissues to promote the formation and growth of the neuromuscular synapses. Moreover, we identified that TDP-43 regulates the expression levels of Disc-large (Dlg) and demonstrated that the modulation of Dlg activity, in skeletal muscles or motoneurons, was sufficient to recover the TDP-43-null locomotive and synaptic defects in flies. Additionally, we found that similar mechanisms are conserved in human cell lines and present in tissues derived from ALS patients.

**Conclusions:** Our results uncover the physiological role of TDP-43 in skeletal muscles as well as the mechanisms behind the autonomous and non-autonomous behavior of this protein in the organization of the neuromuscular synapses.

## BACKGROUND

Amyotrophic lateral sclerosis (ALS) is a devastating disease characterized by the progressive denervation of skeletal muscles followed by motoneurons degeneration and loss. In relation with the pathological origin of the disease, biochemical defects and genetic mutations in the ribonuclear protein (RNP) TDP-43 were correlated with the neurological symptoms of ALS and detected in the great majority of affected patients [1, 2]. Accordingly, histological studies performed in ALS nervous system revealed the presence of protein inclusions made of misfolded, and abnormally phosphorylated, TDP-43 [3–5]. These pathological modifications, affect the subcellular distribution of TDP-43 which abandon its normal localization in the cell nucleus to become redistributed throughout the cytoplasm. Notwithstanding, that the alterations in TDP-43 described above were predominantly identified in upper and lower motoneurons, additional brain regions and cell types were recently implicated in the pathological process. In this direction, the critical role of the glia and immune cells in the progression of the disease has been recently described in ALS individuals and confirmed in different animal models (Boillée et al., 2006; Brettschneider et al., 2012; Diaper et al., 2013; Ince et al., 2011; Tong et al., 2013). Regarding to that, we have reported that the endogenous TDP-43 protein in Drosophila *melanogaster* (TBPH) localizes in glial cells and is required in these tissues to prevent motoneurons degeneration supporting the idea that ALS may present a non-neuronal origin [10]. In consonance with this hypothesis, pathological modifications in TDP-43 were also detected in the skeletal muscles of patients suffering of different neuromuscular diseases alike inclusion body myositis (IBM) or sporadic and familial forms of ALS [12]. In the same direction, overexpression of TDP-43 led to age-related muscles weakness and degeneration in mice [13], zebrafish [14] and Drosophila [15, 16], suggesting that the tight regulation of TDP-43 function may play an important role in muscles physiology. Additionally, it was recently described that TDP-43 is required for the differentiation of C2C22 myoblasts *in vitro* and necessary for the regeneration of tibial muscles *in vivo* [17]. However, the identity of the molecules and metabolic pathways regulated by TDP-43 in skeletal muscles remain largely unknown. Similarly, the observations described above do not specify whether primary defects in the muscular activity of TDP-43 are able to initiate the degenerative course of the disease *in vivo* or the potential consequences of these alterations in the establishment of the neuromuscular synapses. In order to address these questions, we decided to analyze the endogenous function of the TDP-43 ortholog gene, TBPH, in the skeletal muscles of the fruit fly Drosophila *melanogaster*. As a result, our data reveals that TBPH is required in muscular tissues to promote the formation of the neuromuscular synapses and prevent progressive paralysis and early death.

## RESULTS

### The suppression of TBPH in muscles affects Drosophila locomotion and life span

Regarding to these experiments, we have previously described that TBPH becomes expressed in Drosophila muscles and is present in myocytes form larval stages till adulthood [16]. Therefore, to analyze the role of TBPH in these tissues we decide to suppress the expression of the protein by using RNA interference (RNAi). For these experiments, we utilized two different GAL4 lines: the myosin heavy chain (*Mhc*-GAL4) that predominantly gives expression in larval muscles and the transcription factor Mef2 (*Mef2*-GAL4) who expresses during muscles development and adulthood [18]. As a result, we observed that the expression of an RNAi against TBPH (TBi) with *Mhc*-GAL4 or *Mef2*-GAL4 significantly affected the locomotor capacities of the treated flies at both larval and adult stages (Fig. 1a and b). Moreover, we found that the life span of the *Mef2*-GAL4, TBi expressing flies was strongly compromised compared to the *Mef2*-GAL4, GFP-RNAi expressing controls (GFPi) (Fig. 1c). In addition, we observed that the reduction of one copy of the endogenous gene was able to enhance the locomotive defects occasioned by the expression of TBi (*Mhc*-GAL4, TBPH^-^/_+_, TBi vs *Mhc*-GAL4, TBPH^-^/_+_, GFP-RNAi) indicating that these phenotypes were gene dose sensitive and, therefore, rather specific (Fig. 1a). At the cellular level, we found that the expression of TBi strongly affected the organization of the neuromuscular synapses including the localization of the postsynaptic proteins present in the muscular membranes (Fig. 1d-g). More specifically, we observed that the intensity levels of the postsynaptic protein Disc-large (Dlg) were strongly reduced in TBPH-RNAi treated muscles compared to controls with the presence of numerous, pathological, gaps in the distribution of Dlg around the motoneurons axons (Fig. 1d and e). Furthermore, we detected that the postsynaptic distribution and clustering of the glutamate receptors (GluRIIA) were impaired in the RNAi treated flies related to controls indicating that the muscular function of TBPH is required to prevent the disorganization of the postsynaptic membranes (Fig. 1f and g). Furthermore, we noticed that the downregulation of TBPH in Drosophila muscles induced non-autonomous alterations in the structure of the presynaptic terminals with the loss of the characteristic, round and smooth, shape of the synaptic buttons (Fig. 1h and i). At the molecular level, we found that these morphological defects were associated with the significative reduction in the intensity levels of the presynaptic microtubule binding protein *futsch*, homolog to the human protein MAP1B, and responsible for the organization of the synaptic microtubule cytoskeleton (Fig. 1j and k), suggesting that the postsynaptic function of TBPH might be mainly required for the maintenance of these structures or to prevent muscles denervation.

**Fig. 1.**
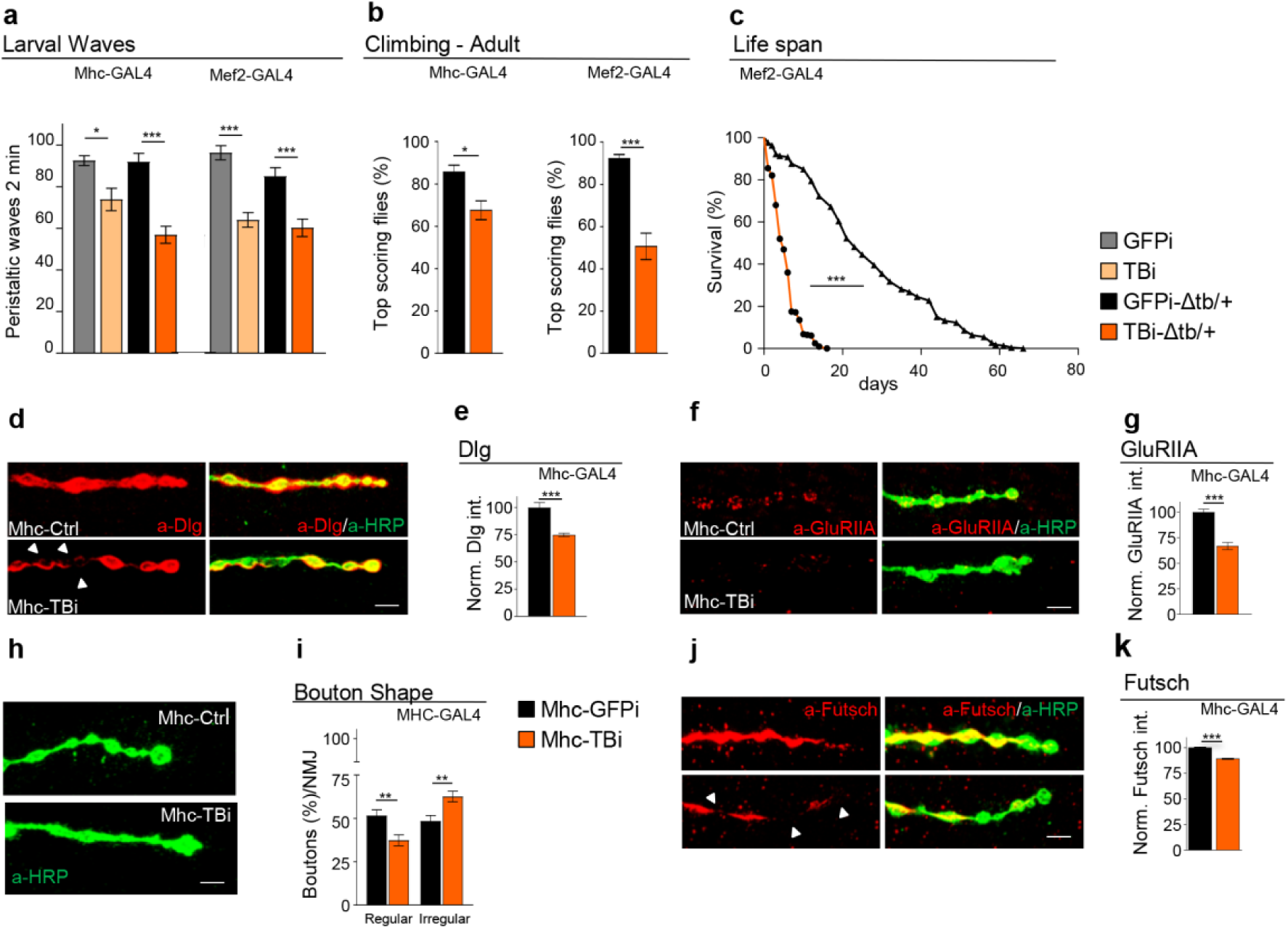
Suppression of TBPH in muscle affects locomotion and life span. **a** Number of peristaltic waves of GFPi (+/+;driverGAL4/UAS-GFP-IR), TBi (+/+;driverGAL4/UAS-TBPH-IR), GFPi-Δtb/+ (tbph^Δ23^/ +;driverGAL4/UAS-GFP-IR) and TBi-Δtb/+ (tbph^Δ23^/+;driverGAL4/UAS-TBPH-IR) using Mhc-GAL4 (left panel) and Mef2-GAL4 (right panel). *n*=20. **b** Climbing assay of GFPi-Δtb/+ and TBi-Δtb/+ using Mhc-GAL4 (left panel) and Mef2-GAL4 (right panel) at day 7. *n*=200. **c** Life span analysis of TBi-Δtb/+ compared to GFPi-Δtb/+ using Mef2-GAL4. *n*=200. **d** Confocal images of third instar NMJ terminals in muscle 6/7 second segment stained with anti-HRP (in green) and anti-Dlg (in red) in Mhc-GFPi (tbph^Δ23^/ +;Mhc-GAL4/UAS-GFP-IR) and Mhc-TBi (tbph^Δ23^/+;Mhc-GAL4/UAS-TBPH-IR). **e** Quantification of Dlg intensity normalized on ctrl. *n*>200 boutons. **f** Confocal images of third instar NMJ terminals in muscle 6/7 second segment stained with anti-HRP (in green) and anti-GluRIIA (in red) in Mhc-GFPi and Mhc-TBi. **g** Quantification of GluRIIA intensity normalized on ctrl. *n*>200 boutons. **h** Confocal images of third instar NMJ terminals in muscle 6/7 second segment stained with anti-HRP (in green) in Mhc-GFPi and Mhc-TBi. **i** Quantification of boutons shape. *n*=200. **j** Confocal images of third instar NMJ terminals in muscle 6/7 second segment stained with anti-HRP (in green) and anti-Futsch (in red) in Mhc-GFPi and Mhc-TBi. **k** Quantification of Futsch intensity normalized on ctrl. *n*>200 boutons. ns= not significant, *p<0.05, **p<0.01, ***p<0.001 calculated by one-way ANOVA (for more than two groups) and T-test (for two groups), log rank test (for survival analysis) error bars SEM. Scale bar 5µm (in d,f, h and j).

### The muscular function of TBPH is sufficient to promote neuromuscular synapses growth and innervation

In order to further characterize the function of TBPH in skeletal muscles, we found that the genetic expression of the endogenous TBPH protein in TBPH null backgrounds (Δtb-TBPH) using the drivers *Mhc*-GAL4 or *Mef2*-GAL4, was sufficient to recover the normal locomotive behavior in larvae and adult flies (Fig. 2a-c and Additional file 1 Fig S1). Similarly, we found that the muscular expression of the human TDP-43 protein in TBPH-null backgrounds (Δtb-hTDP-43) was able to recover Drosophila motility suggesting that the role of these proteins in skeletal muscles must be conserved (Fig. 2a). On the contrary, we found that the expression of the RNA-binding deficient isoform of TBPH (TBPH^F/L^) was not able to recover the TBPH-minus phenotypes demonstrating that the RNA binding ability of the protein is essential. Molecularly, we detected that the recovery of the TBPH function in the muscles of the TBPH minus flies, was sufficient to stimulate motoneurons axons growth, the formation of terminal branches and the addition of new synaptic buttons (Fig. 2d-f). Interestingly, we observed that the non-autonomous rescue of the presynaptic terminals was followed by the reestablishment of the evoked junction potentials (EJPs) between the motoneurons and the underlying muscles suggesting that synaptic transmission was also recovered in TBPH expressing muscles compared to controls (Fig. 2g). Finally, we identified that the expression of the TBPH protein in the muscles of the TBPH-null flies was able to almost completely reestablish the cytoplasmic levels and the postsynaptic distribution of Dlg around the presynaptic terminals (Fig. 2h and i). Moreover, the results described above included the significative reorganization of the glutamate receptors in well-defined clusters at the postsynaptic membranes (Fig. 2j and k).

**Fig. 2.**
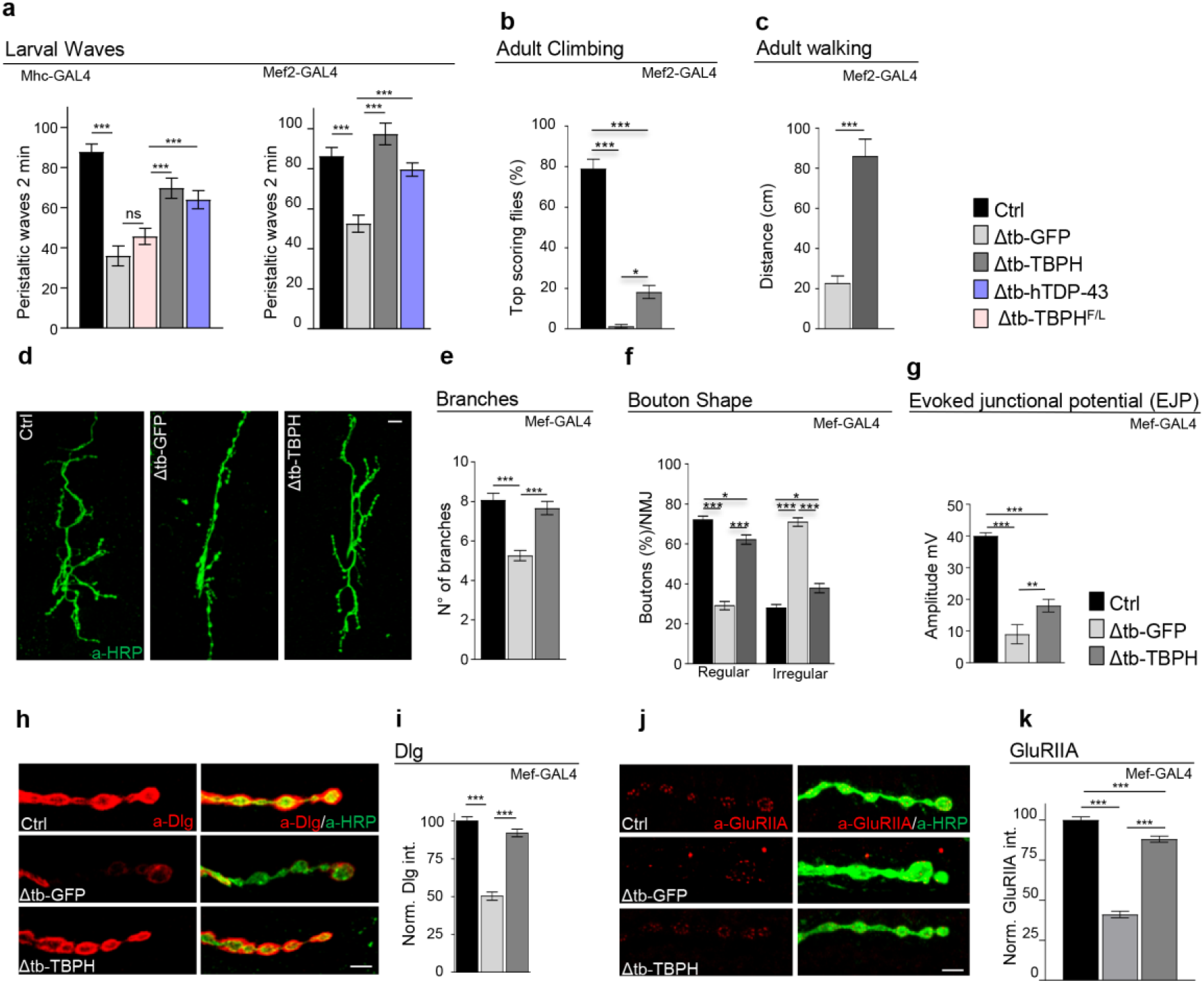
The muscular function of TBPH is sufficient to promote neuromuscular synaptic growth. **a** Number of peristaltic waves of Ctrl (*w*^1118^), Δtb-GFP (tbph^Δ23^/tbph^Δ23^;driver-GAL4/UAS-GFP), Δtb-TBPH (tbph^Δ23^,UAS-TBPH/tbph^Δ23^;driver-GAL4/+), Δtb-hTDP-43 (tbph^Δ23^/tbph^Δ23^;driver-GAL4/UAS-TDP-43) and Δtb-TBPH^F/L^ (tbph^Δ23^/tbph^Δ23^;driver-GAL4/UAS-TBPH^F/L^) using Mhc-GAL4 (left panel) and Mef2-GAL4 (right panel). *n*=20. **b** Climbing assay of Ctrl, Δtb-GFP and Δtb-TBPH using Mef2-GAL4 at day 2. *n*=200. **c** Walking assay analysis of Δtb-GFP and Δtb-TBPH using Mef2-GAL4 at day 2. *n*=100. **d** Confocal images of third instar NMJ terminals in muscle 6/7 second segment stained with anti-HRP (in green) in Ctrl, Δtb-GFP and Δtb-TBPH. **e** Quantification of branches number. *n*=15. **f** Quantification of boutons shape. *n*=200. **g** Evoked neurotransmitter release. Representative EJPs evoked by segmental nerve stimulation of Ctrl, Δtb-GFP and Δtb-TBPH in muscle fiber 6/7 of A3 in third instar larvae. For each fiber 15 EPPs following 0.5 Hz stimulation were considered. **h** Confocal images of third instar NMJ terminals in muscle 6/7 second segment stained with anti-HRP (in green) and anti-Dlg (in red) in Ctrl, Δtb-GFP and Δtb-TBPH. **i** Quantification of Dlg intensity normalized on ctrl. *n*>200 boutons. **j** Confocal images of third instar NMJ terminals in muscle 6/7 second segment stained with anti-HRP (in green) and anti-GluRIIA (in red) in Ctrl, Δtb-GFP and Δtb-TBPH **k** Quantification of GluRIIA intensity normalized on ctrl. *n*>200 boutons. *p<0.05, **p<0.01, ***p<0.001 calculated by one-way ANOVA, error bars SEM. Scale bar 10µm (in d) and 5µm (in h and j).

### TBPH promotes the assembly of the neuromuscular synapses through the regulation of the muscular and neuronal levels of Dlg

Concerning the molecular mechanisms behind the function of TBPH in Drosophila muscles, we have described that the modulation of TBPH levels in these tissues affected the postsynaptic distribution and levels of Dlg (Fig. 1d and e and Fig. 2h and i) suggesting that this protein may contribute to TBPH activity. In agreement with this hypothesis, we found that the mRNA sequence of Dlg presents a series of TBPH putative binding sites that suggest these molecules could physically interact. To test these possibilities, we expressed a flag tagged version of TBPH in Drosophila muscles to perform RIP assays *in vivo*. Interestingly, we found that TBPH was able to bind the mRNA of Dlg in pull down experiments compared to TBPH^F/L^ (Fig. 3a). In addition, we established that the ectopic expression of Dlg in Drosophila muscles with *Mef2*-GAL4, was sufficient to recover the motility problems and climbing defects described in TBPH null flies (Fig. 3b and c). and a reestablishment of the evoked junction potentials (EJPs) between the motoneurons and the underlying muscles (Fig. 3d). Besides these results, we found that the muscular expression of Dlg was able to significantly recover the postsynaptic organization of the glutamate receptors at the postsynaptic, neuromuscular, membranes (Fig. 3i and j). Surprisingly, we observed that the postsynaptic expression of Dlg was capable to promote the non-autonomous growth of motoneurons axons indicating that Dlg is as a *bona fide* mediator of the signaling pathways generated or regulated by TBPH in Drosophila muscles (Fig. 3e-h). Regarding to these results, the presence of Dlg expression was also detected in motoneurons axons suggesting that TBPH might have a similar role in the regulation of Dlg expression in neuronal tissues. In support of this idea, we found that TBPH was able to bind the mRNA of Dlg in RIP assays performed in Drosophila brains after the expression of TBPH with the pan-neuronal driver *elav-*GAL4 (Fig. 4a). Moreover, we observed that Dlg appeared downregulated in adult brains of Drosophila TBPH^Δ23^ and TBPH^Δ142^ homozygous alleles compared to wildtype controls (Fig. 4b). Additionally, we detected that the defects in Dlg levels were recuperated following the genetic expression of the endogenous TBPH protein demonstrating that these differences were rather specific. At the functional level, we identified that the presynaptic expression of Dlg in TBPH-minus flies promoted the recovery of the locomotive problems described in TBPH-null larvae (Fig. 4c) and stimulated the presynaptic growth of motoneurons axons with the formation of new terminal branches and synaptic boutons (Fig. 4d and e). Furthermore, we noticed that the neuronal expression of Dlg induced the non-autonomous clustering of the glutamate receptors present in the postsynaptic membranes of the TBPH-minus NMJs (Fig. 4f and g). Altogether, the data demonstrates that TBPH regulates the pre-and postsynaptic levels of Dlg and its activity is required in muscles and neurons to promote the formation of the NMJs.

**Fig. 3.**
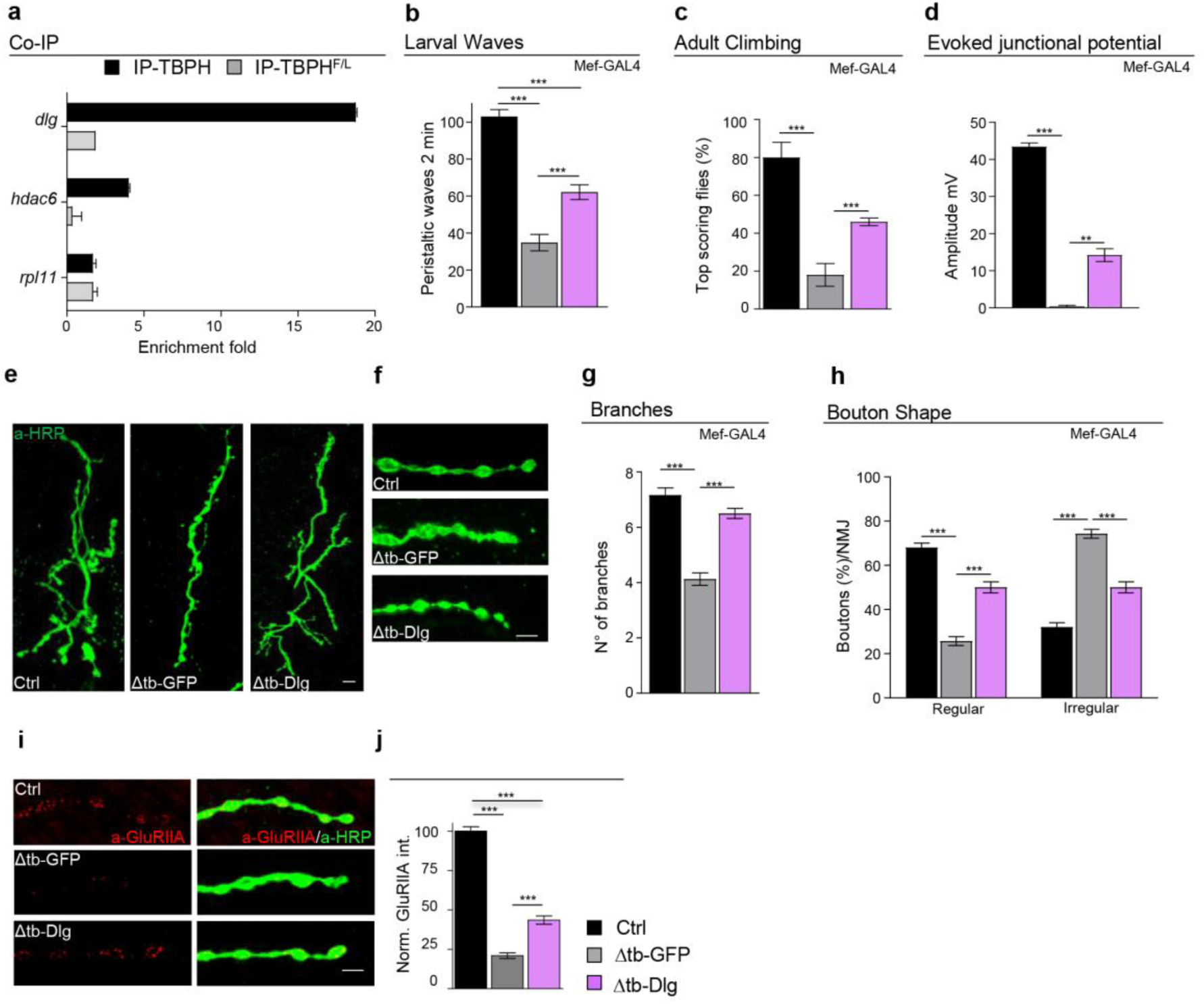
TBPH in muscle promotes synaptic growth through the regulation of Dlg levels. **a** qRT-PCR analysis of mRNAs immunoprecipitated by Flag-tagged TBPH (UAS-TBPH/+;Mef2-GAL4/+, IP-TBPH) and its mutant variants TBPH^F/L^ (+/+;UAS-TBPH ^F/L^/Mef2-GAL4, IP-TBPH^F/L^) in adult thoraxes. The *dlg* enrichment-folds was referred to *rpl-11* (negative control), *hdac6* has been used as positive control. *n*=3 (biological replicates). **b** Number of peristaltic waves of Ctrl (*w*^1118^), Δtb-GFP (tbph^Δ23^/tbph^Δ23^;Mef2-GAL4/UAS-GFP) and Δtb-Dlg (tbph^Δ23^,UAS-Dlg/tbph^Δ23^;Mef2-GAL4/+). *n*=20. **c** Climbing assay of Ctrl, Δtb-GFP and Δtb-Dlg using Mef2-GAL4 at day 2. *n*=200. **d** Evoked neurotransmitter release. Representative EJPs evoked by segmental nerve stimulation of Ctrl, Δtb-GFP and Δtb-Dlg in muscle fiber 6/7 of A3 in third instar larvae. For each fiber 15 EPPs following 0.5 Hz stimulation were considered. **e** Confocal images of third instar NMJ terminals in muscle 6/7 second segment stained with anti-HRP (in green) in Ctrl, Δtb-GFP and Δtb-Dlg. **f** Confocal images of third instar NMJ terminal boutons in muscle 6/7 second segment stained with anti-HRP (in green) in Ctrl, Δtb-GFP and Δtb-Dlg. **g** Quantification of branches number. *n*=15. **h** Quantification of boutons shape. *n*=200. **i** Confocal images of third instar NMJ terminals in muscle 6/7 second segment stained with anti-HRP (in green) and anti-GluRIIA (in red) in Ctrl, Δtb-GFP and Δtb-Dlg. **j** Quantification of GluRIIA intensity normalized on ctrl. *n*>200 boutons. **p<0.01, ***p<0.001 calculated by one-way ANOVA, error bars SEM. Scale bar 10µm (in e) and 5µm (in f and i).

**Fig. 4.**
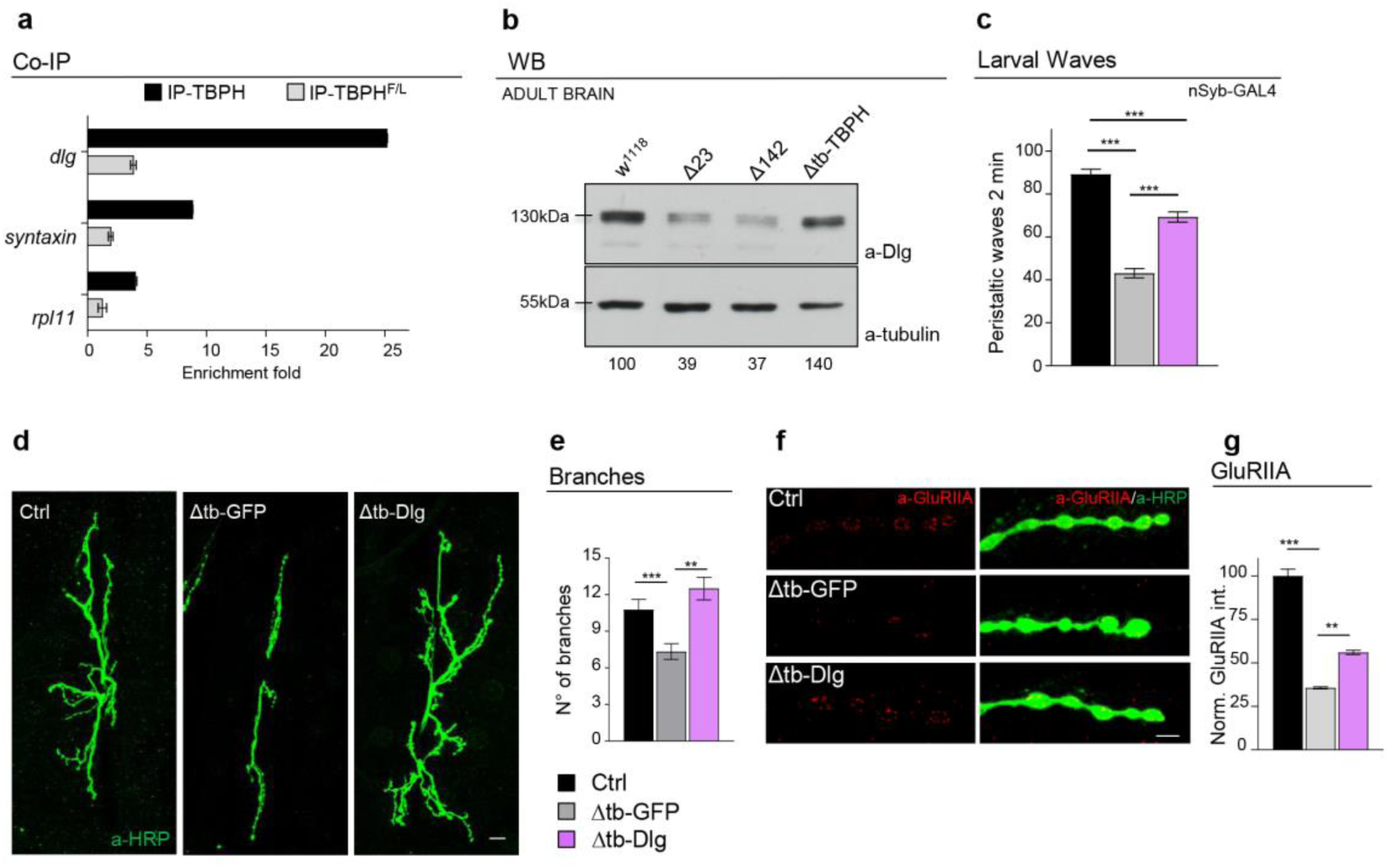
TBPH in neurons promotes synaptic growth through the regulation of Dlg levels. **a** qRT-PCR analysis of mRNAs immunoprecipitated by Flag-tagged TBPH (Elav-GAL4/UAS-TBPH/+;+/+, IP-TBPH) and its mutant variants TBPH^F/L^ (Elav-GAL4/+;UAS-TBPH ^F/L^/+, IP-TBPH^F/L^) in adult heads. The *dlg* enrichment-folds was referred to *rpl-11* (negative control), *syntaxin* has been used as positive control. *n*=3 (biological replicates). **b** Western blot analysis of lane 1 (*w*^1118^), lane 2 (tbph^Δ23^/tbph^Δ23^), lane 3 (tbph^Δ142^/tbph^Δ142^) and lane 4 (tbph^Δ23^,Elav-GAL4/tbph^Δ23^,UAS-TBPH). Adult brains, 1 day old, were probed with anti-Dlg and alpha-tubulin antibodies. The same membrane was probe with the two antibodies and the bands of interest were cropped. Quantification of normalized amounts was reported below each lane. *n*=3 (biological replicates). **c** Number of peristaltic waves of Ctrl (*w*^1118^), Δtb-GFP (tbph^Δ23^/tbph^Δ23^;nSyb-GAL4/UAS-GFP) and Δtb-Dlg (tbph^Δ23^,UAS-Dlg/tbph^Δ23^;nSyb-GAL4/). *n*=20. **d** Confocal images of third instar NMJ terminals in muscle 6/7 second segment stained with anti-HRP (in green) in Ctrl, Δtb-GFP and Δtb-Dlg. **e** Quantification of branches number. *n*=15. **f** Confocal images of third instar NMJ terminals in muscle 6/7 second segment stained with anti-HRP (in green) and anti-GluRIIA (in red) in Ctrl, Δtb-GFP and Δtb-Dlg. **g** Quantification of GluRIIA intensity normalized on ctrl. *n*>200 boutons. **p<0.01, ***p<0.001 calculated by one-way ANOVA, error bars SEM. Scale bar 10µm (in d) and 5µm (in f).

### Defects in Dlg levels are conserved in human cells and motoneurons derived from ALS patients carrying mutations in TDP-43

In order to explore whether analogous defects in Dlg levels (induced by the suppression of TBPH) are conserved in human cells, we treated SHSY5Y cells with an RNAi against TDP-43. Consequently, we found that the suppression of TDP-43 provoked the downregulation of the Dlg homolog protein PSD-95 *in vitro*, indicating that these regulatory mechanisms might be present and conserved in human tissues (Fig. 5a). Regarding to these observations, we noticed that motoneurons differentiated from iPS cell lines obtained from patients containing mutations in TDP-43, presented similar reductions in the expression levels of PSD-95 compared to healthy controls (Fig. 5b) suggesting that analogous mechanisms could be implicated in the pathogenesis of ALS.

**Fig. 5.**
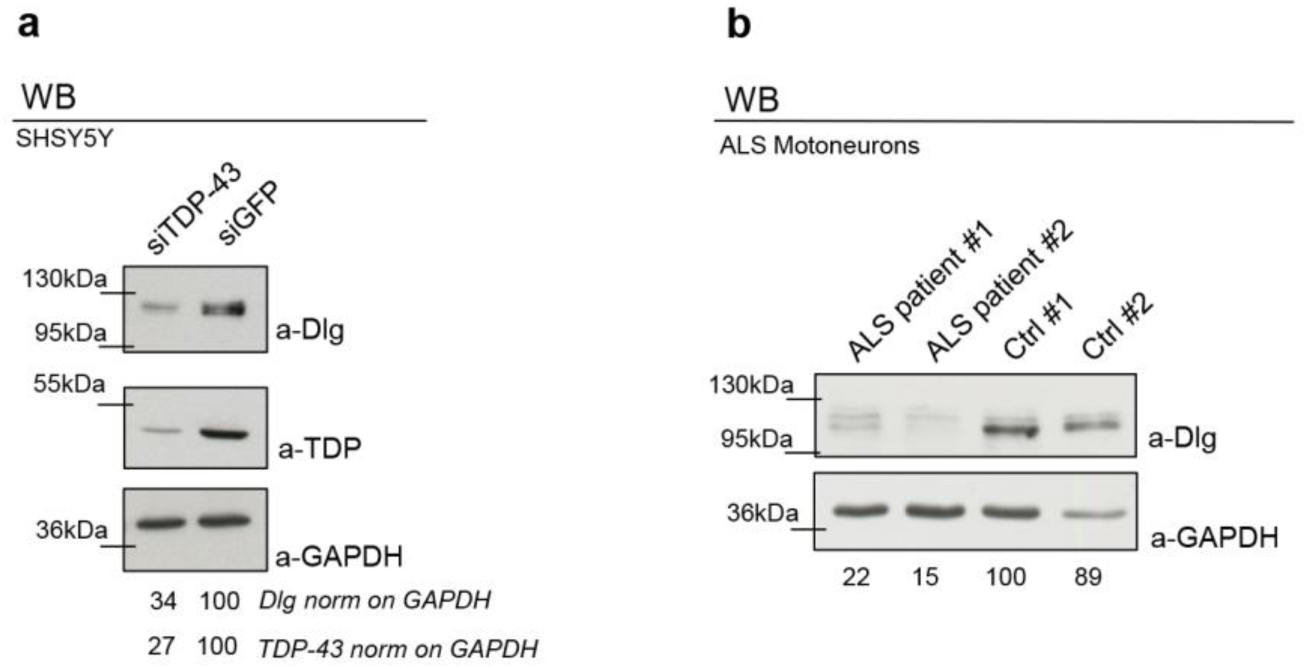
Dlg defects are conserved in human cells and motoneurons derived from ALS patients carrying TDP-43 mutations. **a** Western blot analysis on human neuroblastoma (SH-S5Y5) cell line probed for anti-Dlg, anti-GAPDH and anti-TDP-43 in siGFP (GFP ctrl) and siTDP-43 (TDP-43 silenced). The same membrane was probe with the three antibodies and the bands of interest were cropped. Quantification of normalized protein amount was reported below each lane, *n*=3 (biological replicates). **b** Western blot analysis probed for anti-Dlg and anti-GAPDH on human differentiated motoneurons derived from iPSCs of an ALS patient (ALS patient #1 and ALS patient #2) and a healthy control (Ctrl #1 and Ctrl #2). The same membrane was probe with the two antibodies and the bands of interest were cropped. Quantification of normalized protein amount was reported below each lane.

## DISCUSSION

The atrophy and weaknesses of the body muscles is an essential characteristic of ALS that has been predominantly attributed to motoneurons degeneration. Still, studies performed in patients and experimental systems are providing indications that muscles may play a primary role in the beginnings of the disease. In this regard, we described that the muscular suppression of TBPH provoked locomotive defects with paralysis and a strong reduction in the life span. At the molecular levels, we found that TBPH was required to preserve the postsynaptic organization of the neuromuscular junctions that include the subcellular distribution of Dlg and the GluRIIA at the muscular membranes (Fig. 1d-g). In addition, we did not observe evident signs of muscles degeneration or atrophy after the expression of TBPH RNAi during larval development. In agreement with these results, we did not find alterations during skeletal muscles development in TBPH-null alleles, suggesting that the role of TBPH in Drosophila muscles might be restricted to the formation and differentiation of the neuromuscular synapses [19].

### TBPH supports axonal grow and postsynaptic differentiation in Drosophila muscles and is required to prevent motoneurons axons degeneration

Regarding to the evidences described above, we uncovered that the rescue of the muscular function of TBPH was sufficient to recover the molecular organization of the postsynaptic terminals, including the wild-type sharing of Dlg and GluRIIA. Moreover, we found that the muscular expression of TBPH induced the non-autonomous grow of the motoneurons axonal terminals and synaptic transmission in TBPH-minus flies, followed by the recovery of the locomotive behaviors and life span. Besides, we described that the neurotrophic characteristics of TBPH depended on the RNA-binding capacities of this protein and were conserved in the human ortholog TDP-43 suggesting that similar alterations can be expected in patients with functional defects in TDP-43.

### TBPH regulates the muscular and neuronal levels of Dlg to promote the formation of the neuromuscular synapses

Regarding to the issues mentioned above, in this manuscript we described that TBPH regulates the expression levels of Dlg through direct interactions with its mRNA. Moreover, we demonstrated that these molecular interactions and regulatory mechanisms are present in skeletal muscles and motoneurons. In addition, we showed that the genetic rescue of the Dlg expression levels in the pre- or the postsynaptic compartments was sufficient to recover the neurological defects occasioned by the absence of TBPH. Regarding to that, the capacity of Dlg to recruit scaffolding and signaling proteins, through its PDZ domains, to the plasma membranes was previously described and, would explain the autonomous and non-autonomous roles of TBPH in the formation of the synaptic terminals [20]. In this context, the molecular partners of Dlg, like the adhesion protein *fasciclin* II, might play an important role in the synaptic-organizing functions mediated by TBPH as well as in the mechanisms of ALS. Nevertheless, further experiments would be necessary to verify these hypotheses.

In conclusion, our studies show that primary defects in TBPH function in skeletal muscles provoke locomotive problems and affect Drosophila life span. Molecularly, we describe that TBPH binds the mRNA of Dlg and regulates the expression levels of this protein in muscles and/or motoneurons to regulate the assembly and functional organization of the neuromuscular junctions. In addition, we found that these mechanisms are conserved in human cell lines and present in tissues derived from patients with ALS.

## METHODS

### Fly strains and maintenance

The following genotypes were used:

*w*^1118^ - w;tbph^Δ23^/CyO^GFP^ - w;tbph^Δ142^/CyO^GFP^ – w;; Mef2-GAL4/TM3,Sb – w;;Mhc- GAL4/TM3,Sb – w;UAS-mCD8::GFP/CyO – w;UAS-TBPH/CyO – w;;UAS- TBPH^F/L^/TM3,Sb – w;;UAS-hTDP/TM3,Sb - w;UAS-DLG^EGFP^ - UAS-TBPH-RNAi (#ID38377) - UAS-GFP-RNAi (#9330) - UAS-Dcr-2.

### Larval movement

Wandering 3^rd^ instar larvae were picked from tubes and washed in a drop of demineralized water. A single larva at the time was transferred into a 10 cm diameter dish, filled with 0,7% agar. After 30 s of adaption period, the number of peristaltic waves were counted for a period of 2 min. Generally, 20 – 25 larvae per genotype were tested.

### Survival rate

1 day old adult flies were collected from the fly tube of experimental cross in a 1:1 proportion of female and male and transferred to fresh fly tube and stored in the incubator under controlled conditions (25°C and 60% humidity, 12 h light and 12 h night). Every second day flies were transferred into a fresh fly tube without anesthesia and the number of deaths was scored. A minimum of 200 flies per genotype were tested.

### Climbing assay

1 day old adult flies were collected from the fly tube of experimental cross in a 1:1 proportion of female and male and transferred into fresh fly tube and maintained in an incubator as previously described. A 50 ml glass cylinder was divided into three parts, as bottom, middle and top (5 cm each part). Flies were carefully flipped into the cylinder from the fly tube without any anesthesia and gently dropped to the bottom. After 30 s of adaptation period, flies were dropped again onto the bottom of the cylinder and after the time interval of 15 s the numbers of flies present in each part of the cylinder were scored. For each genotype 3 trials per each tube were done and the average of the scored fly numbers was considered as the final score. A minimum of 200 flies was tested for each genotype.

### Walking assay

Young flies 2-3 days old were tested for walking ability. A 145 mm dish was used. The bottom surface was divided in a grid of 1cm × 1cm squares to facilitate the measuring of the distance walked by flies. The fly without any anesthesia was placed in the middle of the dish and after 30 s of adaptation to the environment the distance walked by the fly was recorded for 30 s, counting the number of squares. A minimum of 50 flies were individually tested for each genotype.

### Immunohistchemistry

Wandering 3^rd^ instar larvae were picked from fly tube, in a drop of demineralized water and selected for the genotype. Individually picked larva was dissected on Sylgard plates, in Dissection Solution (128 mM NaCl, 2 mM KCl, 4 mM MgCl_2_, 0,1 mM CaCl_2_, 35,5 mM Sucrose and 5 mM Hepes (pH 7,2)). Larvae were pinned at both ends with minute pins (Austerlic Isect Pins 0,1 mm diameter, Fine Science Tools, Germany) and opened on the dorsal site with Spring scissors (Fine Science Tools, Germany). Once larva was opened, internal organs were removed and the interior was extensively washed with Dissection Solution leaving muscle wall opened, pinned flat on the surface. The subsequent step was a fixation, generally done with 4% PFA in PBS for 20 min, however, in the case of glutamate receptors staining a methanol fixation of 5 min at −20°C was performed. Fixation solution was removed with 3 washes in PBS-T (PBS 1x supplemented with 0,1% (v/v) Tween20) for 5 min each. After a blocking step of 30 min in blocking solution (5 % NGS (Normal Goat Serum (#S-1000 Chemicon) in PBS-T buffer), larvae were incubated over night at 4°C in primary antibodies diluted in blocking solution. The day after, primary antibody was removed with three washes of 10 min each with PBS-T and a further blocking step of 30 min was performed before secondary antibodies addition. All secondary antibodies were diluted in blocking solution. An incubation was 2 h long, carried out at the room temperature. Excess of antibody was removed by 3 subsequent washes of 20 minutes each in PBS-T. Finally, dissected-stained larvae were incubated over night at 4°C in Slowfade®Gold antifade (#S36936 Life Technologies) reagent, before being mounted on a glass slide. anti-HRP (#323-005-021 Jackson 1:150), anti-GFP (#A11122 Life Technologies 1:200), anti-GluRIIA 8B4D2c (DSHB 1:15), anti-Dlg 4F3c (DSHB 1:250), anti-Futsch 22C10s (DSHB 1:50) Alexa-Fluor® 488 (mouse #A11001 or rabbit #A11008 1:500) and Alexa-Fluor® 555 (mouse #A21422 or rabbit #A21428 1:500).

### Acquisition and quantification of confocal images

In each experiment, the genotypes of interest were processed simultaneously, and the images were acquired using the same settings. Images of muscle 6 and 7 on second abdominal segment were acquired using the LSM Zeiss Software on a Zeiss 510Meta confocal microscope (63 × oil lens) and then analyzed using ImageJ (Wayne Rasband, NIH). For the quantification of pre and postsynaptic markers, samples were double labelled with anti-HRP and the marker of interest: the ratio between the mean intensity of the marker and the HRP was calculated for each bouton of the terminal.

### Electrophysiology on NMJ of the third instar larva preparation

Larval body wall preparations were dissected out in Ca2+-free HL3 solution from third instar larvae pinned on Sylgard coated petri dishes. Central nervous system was excised by cutting segmental nerves roots. After replacing Ca2+ free HL3 solution with Ca2+ 1 mM HL3, post-synaptic potentials at neuromuscular junction of fiber 6/7 of abdominal segments A3/A4 were intracellularly recorded, at room temperature in current-clamp condition, using an intracellular microelectrode (tip diameter 0.5 μm, 15 MΩ resistance). The recorded signal were amplified by a current-clamp amplifier (SEC 05, NPI, Tamm, Germany), digitized at 10 kHz sampling rate using an A/D interface (National Instruments, Austin, TX, USA) and fed to a computer for display and storage using an appropriate software (Win EDR, Strathclyde University, Glasgow, UK).

Fibers with a resting membrane potential below −60 mV were considered for the experiment. In these fibers, membrane potential was set at −70 mV throughout the experiment by injecting current through the intracellular electrode. Evoked postsynaptic potentials (EPSPs or Excitatory Junctional Potentials or EJPs) were recorded by stimulating at 0.1 Hz (pulse duration 0.4 ms; 1.5 threshold voltage) the segmental nerve using a suction electrode (tip diameter ∼10 μm) connected to a stimulator (S88, Grass, Pleasanton, CA, USA) through a stimulus isolation unit (SIU5, Grass, Pleasanton, CA, USA). Intracellular recordings were analyzed offline using pClamp software (pClamp, Axon, Sunnyvale, CA, USA). Statistical comparisons and graphs were made using Graphpad software (Graphpad, La Jolla, CA, USA) or MATLAB (Matworks, Natick, MA, USA).

### Immunoprecipitation

Protein G magnetic beads (#10003D Invitrogen) were washed two times with PBS + 0.02% Tween and coated with anti-FLAG M2 monoclonal antibody (#F3165 Sigma). Thoraces or heads of adult flies were cut and stored in lysis buffer containing 20 mM Hepes, 150 mM NaCl, 0.5 mM EDTA, 10% Glycerol, 0.1% Triton X-100, 1 mM DTT and protease inhibitor (#04 693 159 001 Roche). Samples were homogenized with a Dounce homogenizer, and major debris were removed by centrifugation step of 5 min at 0.4xg at 8°C. The pretreated beads and tissue extracts were mixed and incubated for 30 min at 4°C. After this binding step, beads were washed five times with washing buffer (20 mM Hepes, 150 mM NaCl, 0.5 mM EDTA, 10% Glycerol, 0.1% Triton X-100, 1 mM DTT, protease inhibitor, 0,2 % DOC, 0,5 M Urea) using DynaMagTM-Spin (#123.20D Invitrogen). RNA transcripts bound by TBPH-Flag tagged were extracted. The beads were treated with Trizol (#15596026 Ambion) and precipitated with isopropanol adding glycogen (#R0551 Thermo scientific). Retro-transcription with Superscript III First-Strand Synthesis (#18080-093 Invitrogen) was performed with Oligo-dT primers on resuspended RNA. Real Time PCR was carried out with gene specific primers, which sequences are listed below.

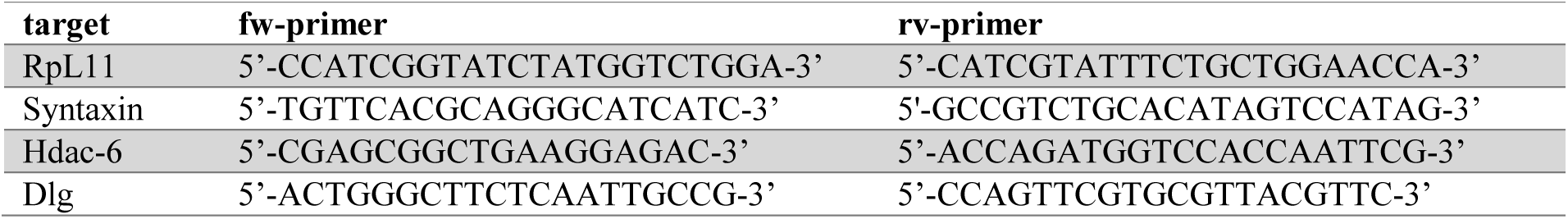

In order to calculate the enrichment fold, initially, all data were normalized to the respective inputs. The signal was represented by how many more fold increase was measured compared to the control signal. The enrichment was calculated according to the following equation:

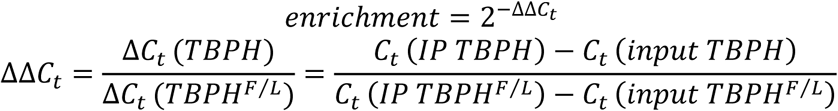

The results were derived from three independent immunoprecipitation experiments.

### Protein extraction

To collect adult heads, flies were flash-frozen in liquid nitrogen for 10 seconds and immediately vortexed to easily detach heads from bodies. Heads were subsequently transferred into Lysis buffer (150 mM Tris, 5 mM EDTA, 10 % glycerol, 5 mM EGTA, 50 mM NaF, 4 M urea, 5 mM DTT and protease inhibitors (#04 693 159 001 Roche). After a squeezing step, performed both, manually and mechanically, the homogenized samples were gotten rid of major debris by centrifugation at 0,5 x g for 6 min on 4°C. The protein concentration of the collected supernatant was quantified with Quant-iT™ Protein Assay Kit (#Q33212 Invitrogen), following supplier protocol.

Transfected neuroblastoma cell line SH-SY-5Y were resuspended in iced RIPA buffer added of protease inhibitors (#04 693 159 001 Roche) and subjected to sonication (Biorupture sonication system, Diagenode).

Lysates were quantified (BCA Protein kit #23225 Thermo Scientific), following supplier protocol.

### SDS-PAGE

Protein samples were diluted in 1x Laemmli buffer (composition of 5x: 0.3 M Tris-HCl pH 6.8, 50 % glycerol, 10 % SDS, 25 % β-mercaptoethanol, 0.05 % bromophenol blue) to reach the same concentration among all and then boiled at 95°C for 5 min. Afterwards they were loaded on a polyacrylamide gel.

The loaded gel was placed into chamber with 1x running buffer (10x running buffer: 30.28 g Tris, 114.13 g Glycine, 10 g SDS in 1 l water). The conditions set were 25 mA per gel.

### Western blot

When proteins were separated by the electrophoresis, they were transferred to a nitrocellulose membrane Amersham™ Protran™ 0.2 µm NC (Life Science). The western blot sandwich was put into the chamber, filled with transfer buffer 1x containing 20 % methanol (transfer buffer 10x: 30 g Tris, 144 g glycine in 1 l water). The transfer lasted 1 hour at 350 mA. The membrane was incubated with a solution of 5 % milk in 1x TBS 0.01 % Tween (TBS-T) for 30 min at a room temperature on a shaker (TBS buffer 10x: 24.2 g Tris, 80 g NaCl in 1 l water, pH 7.6). After blocking, the membrane was set into dilution of primary antibody with TBS-T with 5 % milk. It was placed at 4°C over night. When the incubation with primary antibodiy was over, five washes with TBS-T followed, 5 min each. Next, the membrane was incubated with the secondary antibody diluted in TBS-T with 5 % milk for 1 hour at room temperature. The protein detection was performed with SuperSignal®West Femto Maximum Sensitivity Substrate (#TB260893 Thermo Fisher Scientific). Primary antibodies: anti-Dlg 4F3c (DSHB 1:10000), anti-TBPH (home-made 1:4000), anti-tubulin (#CP06 Calbiochem 1:4000), anti-TDP-43 (#12892-1-AP Proteintech 1:4000), anti-Dlg 2D11 (#SC9961 Santa Cruz 1:1000), anti-GAPDH (#SC25778 Santa Cruz 1:2000). Secondary antibodies: anti-mouse-HRP (#31430 Thermo Fisher Scientific 1:30000(flies), 1:10000(cells)), anti-rabbit-HRP (#31460 Thermo Fisher Scientific 1:10000).

### Cell culture and RNA interference

SH-SY-5Y neuroblastoma cell line was cultured in standard conditions in DMEM-Glutamax (#31966-021, Thermo Fisher Scientific) supplemented 10% fetal bovine serum and 1 × antibiotic-antimycotic solution (#A5955; Sigma). RNA interference of TDP-43 was achieved using HiPerfect Transfection Reagent (#301705, Qiagen) and siRNA specific for human TDP43 (5′-gcaaagccaagaugagccu-3′); as control siRNA for Luciferase was used (5′-uaaggcuaugaagagauac-3′; Sigma). Immediately before transfection 2–4 × 105 cells were seeded in 6-well plates in 1.4 ml of medium containing 10% fetal serum. A volume of 3 μl of each siRNA (40μM solution in water), was added to 91μl of Opti-MEM I reduced serum medium (#51985-026, Thermo Fisher Scientific), incubated 5 minutes at room temperature and subsequently 6 μl of HiPerfect Transfection Reagent were added. The silencing procedure was performed again after 24 and 48 hours.

### Human iPSC Culture and MN differentiation

Human iPSC culture and MN differentiation were already described in (Romano et al., 2018). All the studies performed with human samples were in compliance with the Code of Ethics of the World Medical association (Declaration of Helsinki) and with the national legislation and institutional guidelines. Briefly, fibroblasts from dermal biopsies (Eurobiobank) from ALS patients (TDP-43 mutations #1:G294V; #2:G378S) and controls were reprogrammed into iPSCs with CytoTune-iPS 2.0 Sendai reprogramming Kit (#A16517, ThermoFisher) and differentiated into MNs with the multistep protocol described by Ng (Ng et al., 2015)

### Statistical analysis

All statistical analysis was performed with Prism (GraphPad, USA) version 5.1. One-way ANOVA with Bonferroni correction and t-test with Man-Whitney correction were applied as statistical test. In all figures all the values were presented as the mean and the standard error of the mean (SEM). Statistical significance was portrayed as *p<0.05, **p<0.01, ***p<0.001.

## DECLARATIONS

### Ethics approval and consent to participate

Not applicable

### Consent for publication

Not applicable

### Availability of data and material

The datasets used and/or analysed during the current study are available from the corresponding author on reasonable request.

### Competing interests

The authors declare that they have no competing interests

### Funding

The present work was supported by ARISLA (CHRONOS) and BENEFICENTIA Stiftung.

### Authors’ contributions

NS, GR, CI, RK performed experiments and discussed the data. AM performed electrophysiology analysis, MN provided human iPS cells and the differentiated motoneurons. FF supervised the work, discussed results and wrote the manuscript.

## Acknowledgements

We thank professor Chun-Fang Wu to provide us DLG transgenic flies. The Bloomington Stock Center and Developmental Studies Hybridoma Bank for stocks and reagents.

---

**Additional file 1 Fig. S1.**
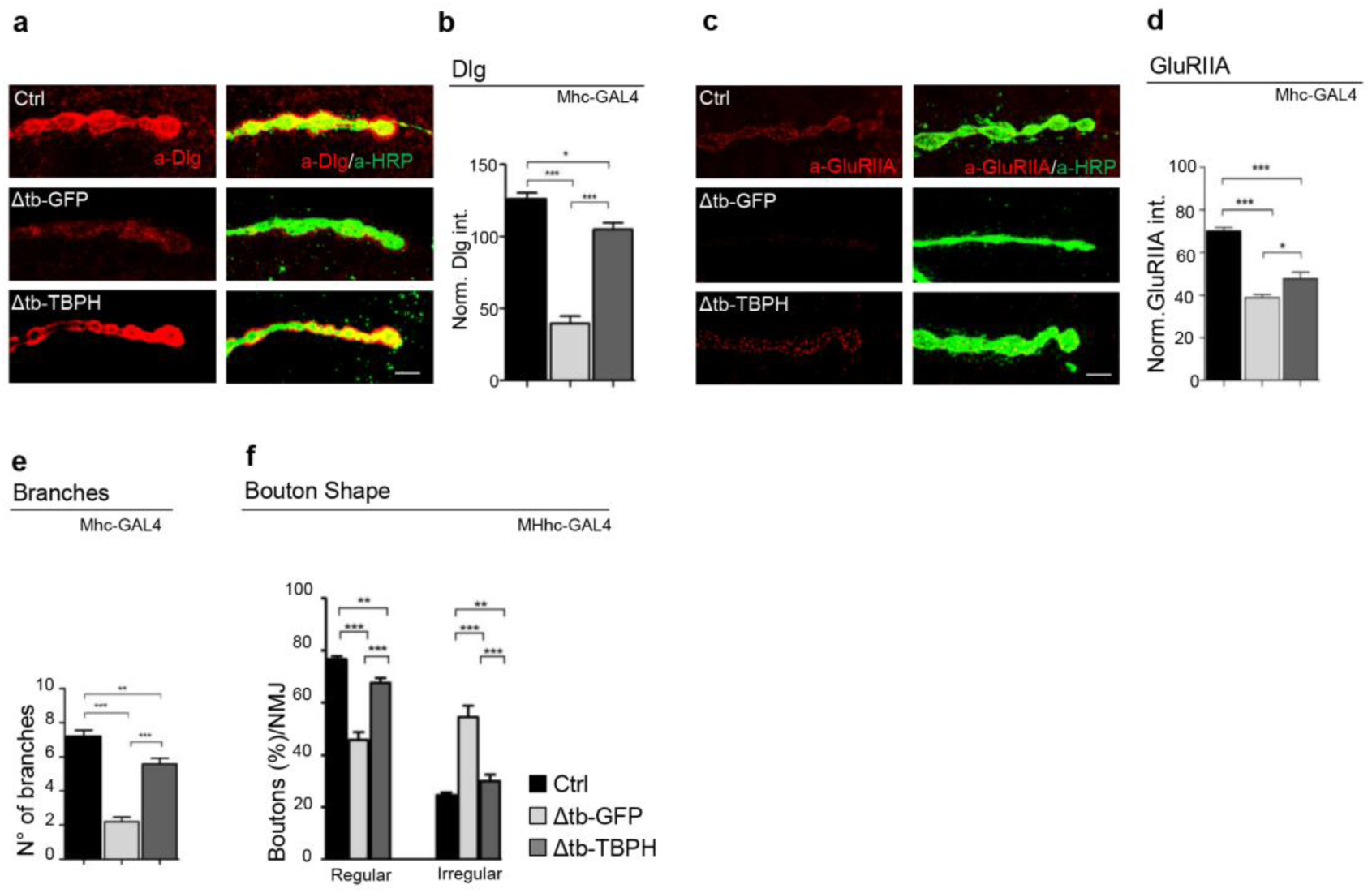
**a** Confocal images of third instar NMJ terminals in muscle 6/7 second segment stained with anti-HRP (in green) and anti-Dlg (in red) in Ctrl (*w*^1118^), Δtb-GFP (tbph^Δ23^/tbph^Δ23^;Mhc-GAL4/UAS-GFP), Δtb-TBPH (tbph^Δ23^,UAS-TBPH/tbph^Δ23^;Mhc-GAL4/+). **b** Quantification of Dlg intensity normalized on ctrl. *n*>200 boutons. **c** Confocal images of third instar NMJ terminals in muscle 6/7 second segment stained with anti-HRP (in green) and anti-GluRIIA (in red) in Ctrl, Δtb-GFP and Δtb-TBPH. **d** Quantification of GluRIIA intensity normalized on ctrl. *n*>200 boutons. **e** Quantification of branches number in Ctrl, Δtb-GFP and Δtb-TBPH. *n*=15. **j** Quantification of boutons shape in Ctrl, Δtb-GFP and Δtb-TBPH. *n*=200.

